# Data-driven identification of the ecologically relevant climatic windows and climate summaries with StatClim model

**DOI:** 10.64898/2026.10.07.757008

**Authors:** Adam Klimeš, Martin Macek, Jan Wild, Josef Brůna, Martin Kopecký

**Affiliations:** Institute of Botany, Czech Academy of Sciences, Zámek 1, 252 43 Průhonice, Czech Republic

**Author notes:** Corresponding author: Adam Klimeš, Zámek 1, 252 43 Průhonice, Czech Republic.

**Keywords:** Bayesian, climwin, lagged effects, microclimate, sliding window, time window

## Abstract

Climate affects biological processes and organisms. It is extensively recorded, and its effects are frequently studied. Climatic variables are recorded as time series of measurements, but these are often summarized as long-term averages, throwing away biologically relevant information. Subsequent analyses can therefore miss important climate effects or suffer from low predictive power.

Here, we propose a new method, StatClim, inferring the most informative climatic summary from the data. Instead of taking a fixed arbitrary summary of a time series, StatClim automatically estimates the beginning and width of a climatic window and simultaneously calculates data aggregation, which best predicts a response variable. We implemented the proposed approach as a Bayesian hierarchical model in Stan.

We illustrated the potential of StatClim on published data on mast seeding of *Fagus sylvatica*. StatClim correctly identified known climatic windows relevant for this tree species, but also uncovered that mast seeding is triggered by high temperatures within these windows. We demonstrated flexibility of the model to account for complex data structures, we ran power analysis showing the ability of the model to identify climatic effects even with modest datasets, and we evaluated its false positive rate, which is lower than 0.01.

We discuss the advantages and limitations of StatClim. Compared to existing alternatives, it is suitable for hypothesis testing and offers important advances, such as estimation of the type of aggregation. In combination with existing methods, StatClim will facilitate data-driven use of climatic and microclimatic data and therefore discovering so far unknown climatic effects on biological processes and ecosystem functioning.

## 1 Introduction

Biological processes respond to climate variability through mechanisms that are temporally structured, often non-linear, and threshold dependent. Consequently, modelling of the relationship between climate and organismal responses requires representations of climate that preserve this temporal complexity rather than reduce it to arbitrary summaries. Climate data is typically available as high-resolution time series of physical variables such as temperature, precipitation, and humidity, but arbitrary long-term averages are often used in ecology and biogeography as predictors of biological responses (Körner and Hiltbrunner 2018). This use of arbitrary summaries inevitably leads to a reduction in information value and weakens the power of ecological models.

Unlike high-resolution climate data, many biological phenomena cannot be assessed with the same frequency and during a comparable timespan due to physical limitations, labor demands, or just the fact that they occur occasionally (e.g., mass reproduction and mortality events). To link climate variability to such phenomena, climatic time series are typically summarized by an average over some pre-defined period (e.g. year). However, this summarization is often arbitrary, and many organisms respond rather to climate extremes (e.g. high temperature, drought events) than to long-term averages (Lenarz et al. 2009; Macek et al. 2019). Similarly, many biological phenomena are driven only by climatic conditions in a specific time window, for example shortly before the phenomenon (e.g., Wyver et al. 2023), in a specific season (e.g., Anderson-Teixeira et al. 2022; Jarecke et al. 2024), or after a certain time (lagged responses; Svenning and Sandel 2013; Evers et al. 2021). As a result, arbitrary climate data summarization throws away the information which could potentially provide insight into the ecology of studied organisms or ecological processes.

To avoid arbitrary summarization of climatic time series, we either need a detailed system knowledge to make the summarization hypothesis-driven (Sollmann 2024), or we need to consider multiple possible summarizations. In ecology, a tested hypothesis often predicts an overall effect of a climatic variable on a phenomenon of interest (e.g., species richness is affected by temperature; Moreno-Rueda and Pizarro 2008) not a specific period for which the climate is relevant, or if the response relies on average or extremes. Currently, the most popular method to approach multiple possible summarizations is model selection using climwin package (also known as sliding window analysis; van de Pol et al., 2016). In brief, climwin first fits many models with different time windows and then selects the best model using the Akaike information criterion (Akaike 1974). While other possible methods exist (Roberts 2008; Brakel 2014), climwin approach generally outperforms these methods (Rubio-Cuadrado et al. 2022; Journé et al. 2025b).

However, model selection might not be suitable for hypothesis testing due to increased probability of false positives (for methods trying to address this issue see: Bailey and van de Pol 2016; van de Pol et al. 2016). Climwin and other existing methods also compare only different time windows and does not include different aggregations within these windows (spanning from minima to maxima). Different aggregations could be theoretically implemented in a model selection approach, but it would often lead to many possible models, making model selection computationally unfeasible. To overcome these limitations, we propose a method which avoids arbitrary summarization, exploits all the climatic variability, including different aggregations, and provides a data-driven assessment of a climate effect while having a low false positive rate. The proposed method, called StatClim, operates within a well-known regression framework and can therefore be combined with other information, for example non-climatic predictors, hierarchical effects, and variables accounting for study design as well as spatial and temporal autocorrelation structures.

Here, we describe the model applied to space-for-time study design (Lovell et al. 2023; Kharouba and Williams 2024), but the model is suitable for a wide range of other design structures (see *Discussion*). In any design, climate data can represent either traditionally used macroclimate or biologically more relevant microclimate, which represent more proximal conditions experienced by the study organisms (Klinges et al. 2024).

## 2 StatClim model

In conventional approaches, climatic predictors are constructed by summarizing time series over predefined periods (e.g., mean temperature over a fixed window of five years). In contrast, StatClim treats this summarization as an estimable component of the model. Rather than specifying the temporal window and aggregation function in advance, both are inferred directly from the data.

StatClim consists of two submodels, which are jointly used to estimate the climatic summary used as a predictor of a biological response:

The first submodel estimates the temporal window used for the summarization by estimating both the starting point and the width of the window. This submodel has the potential to identify, for example, whether precipitation during winter or summer is more relevant for vegetation phenology (Currier and Sala 2022), or determine the period during which the temperature is critical for bird hatching success (Glądalski et al. 2018).

The second submodel estimates the aggregation function within the relevant time window identified by the first submodel. Instead of assuming that biological responses depend on mean conditions, the model evaluates whether responses are driven by specific features of the climatic distribution, such as extremes. For example, this allows testing whether drought events (lower tail of precipitation distribution), rather than average precipitation, determine tree growth (Van Kampen et al. 2022) or whether high temperatures drive the hatching success of birds (McCowan and Griffith 2021).

Because climatic effects are driven by a specific combination of conditions (identified by the second submodel) in a specific window (identified by the first sub-model), both submodels are estimated simultaneously. This joint estimation ensures the correct identification of the climatic summaries even in cases where separated submodels would fail. For example, the importance of summer high temperatures can be identified even if average summer temperatures do not affect the response variable (Fig. 1).

**Figure 1:**
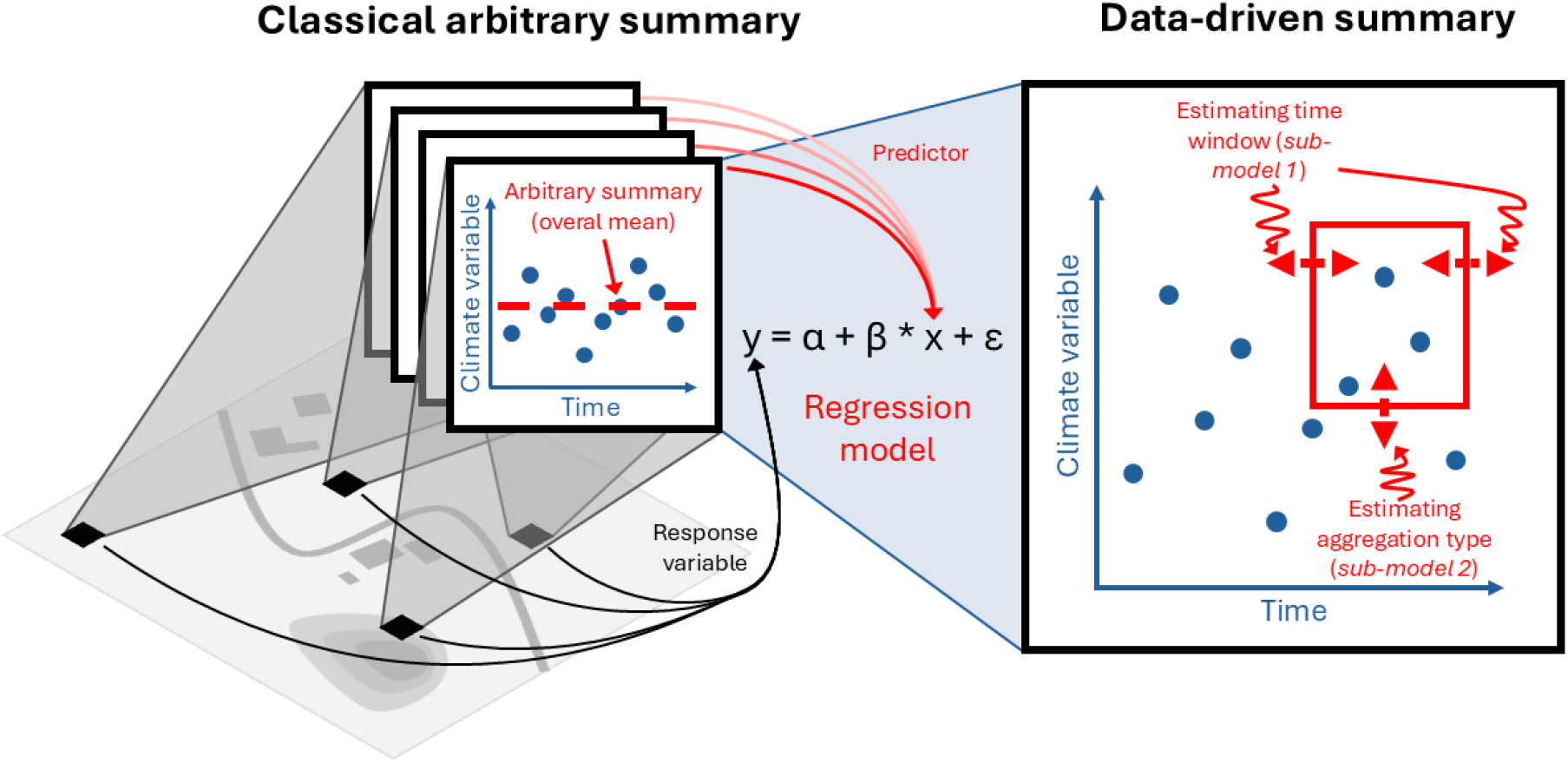
The StatClim model substitutes an arbitrary summarization by data-driven summarization based on three parameters – start of the climatic window, its width, and the type of aggregation.

### 2.1 Implementation

We implemented StatClim as a Bayesian hierarchical model in the probabilistic programming language *Stan* (Carpenter et al. 2017). Both submodels are implemented as logistic window functions determining weights, which are then multiplied and used to calculate a weighted mean. Therefore, the estimated time window and aggregation are not discrete, but there is a biologically realistic smooth transition between values which are used in the summary and values which are not. The window is defined by two parameters controlling the location (starting point) and width (which is fixed for aggregation) and by smoothness of the transition. In ecology, these smooth transitions of the logistic window are more appropriate than hard-edged time windows (e.g., van de Pol and Cockburn 2011). Importantly, logistic window for aggregation is nearly identical to aggregating by percentile, but is differentiable, more flexible, and less sensitive to extreme values (Fig. S1).

To ensure that the aggregating type refers to the specific time window estimated by the first submodel, its logistic window is applied to cumulative weights of sorted values from the first submodel:

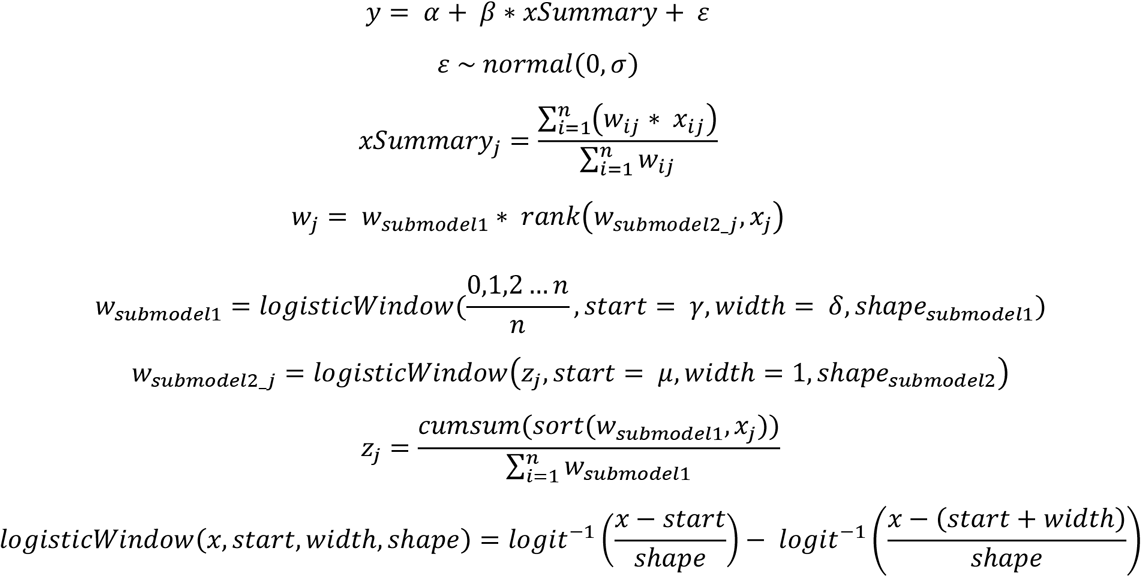

Where α, β are parameters (intercept and slope) of the regression, ‘normal’ stands for normal distribution, σ is a variance parameter, i is the index of observations within a time series (e.g., for individual temperature measurements), j is the index of sites, rank(x, y) is a function that orders vector x based on a rank of values in vector y, γ and δ are parameters of submodel 1 determining the start and width of the logistic window specifying the time window, μ is a parameter of the submodel 2 determining the start of the logistic window specifying the type of aggregation, ‘shape’ are parameters provided by the user which determine the steepness of the logistic windows, cumsum(x) is a function returning the cumulative sum of x, sort(x, y) is a function which sorts vector x based on the order of values in vector y, logit^-1^(x) is inverse logit function.

### 2.2 StatClim outputs

Before demonstrating the potential of the StatClim model (see *Case Study* below), we show outputs of the core model. We ran the model using a simulated dataset of 60 sites each with 100 climate measurements and a response variable. The response variable was simulated to be dependent on high values of the climatic variable in a climatic window extending the first half of the time series using the same dataset, priors, and sampling parameters as for power analysis (see *Power analysis* below). StatClim successfully identified the climatic window at the beginning of the climatic time series and aggregation within this window showing the effect of the highest values of the climatic variable (Fig. 2).

**Figure 2:**
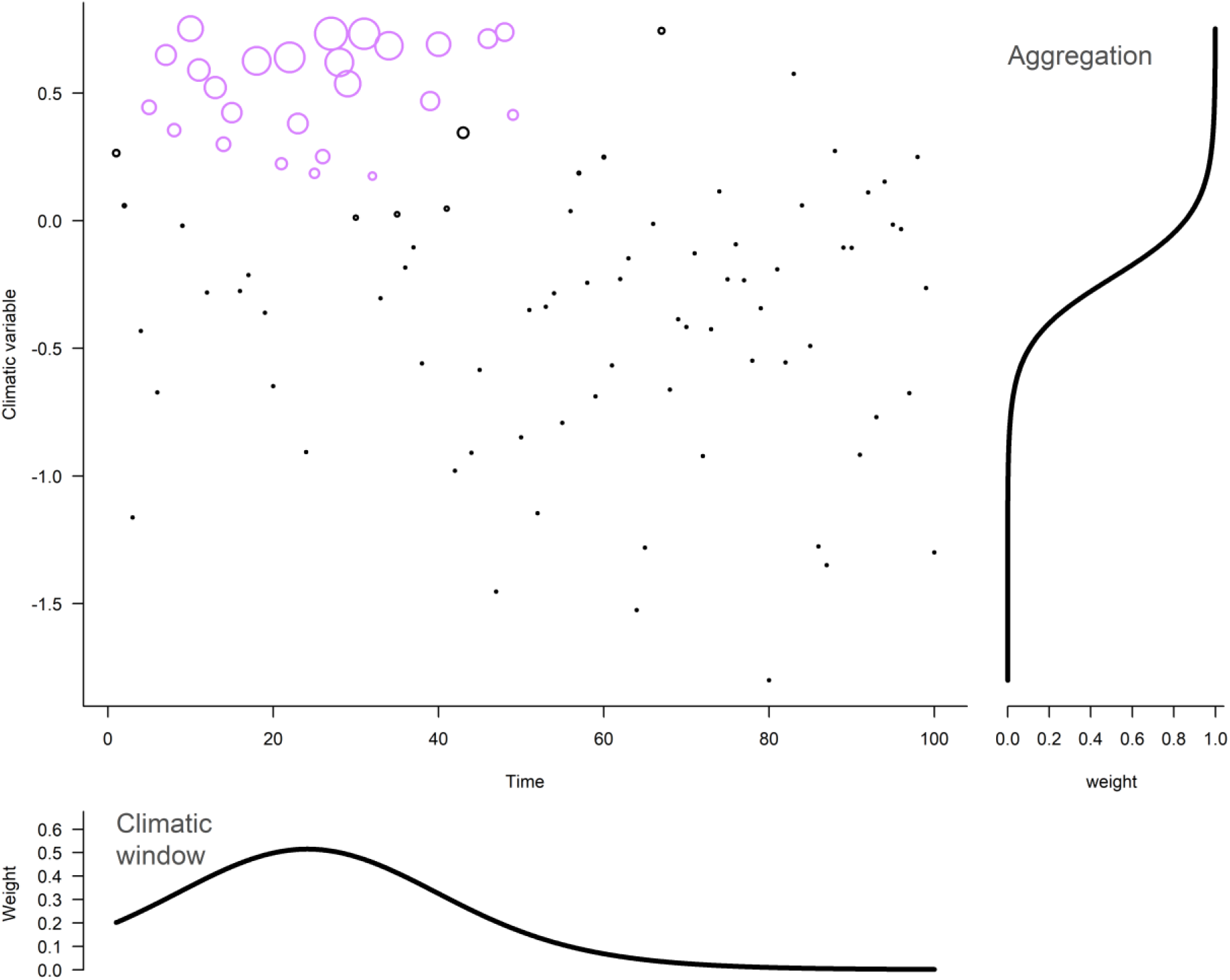
Climatic window and aggregation estimated by the StatClim model (simulated dataset). The scatter plot shows 100 observations of a climatic variable for one site with a point size corresponding to the weight of the observation in the model. From these weights, the weighted mean - summary of the climatic variable for this site was computed and used as a predictor of the response variable. This weight is a product of weights assigned by the climatic window (shown at the bottom) and the aggregation (shown at the right side). Points in purple are observations that were used for response simulation in the data simulation process (true values that were successfully identified using StatClim).

### 2.3 Power analysis

To assess the required sample size for reliable identification of the climatic window and aggregation using StatClim, we ran power analysis in which we simulated datasets with 30, 60, and 90 sites with weak, medium, and strong effect of a climatic variable on a response variable and estimated the effect using StatClim. Specifically, we simulated climate variable time series consisting of 100 values for the given number of sites (30, 60, and 90) and the response variable (one value for each site). The climatic variable was simulated in two parts, one part for the prediction of the response variable representing 25 observations randomly positioned in the first half of the climatic time series and the second part representing the rest with no effect on the response variable. The part with the effect on the response variable had a higher mean value than the rest simulating the effect of climatic extremes. We simulated the response variable with specified strength of the climatic effect (weak, medium, and strong: R^2^ = 0.2, 0.5, and 0.8).

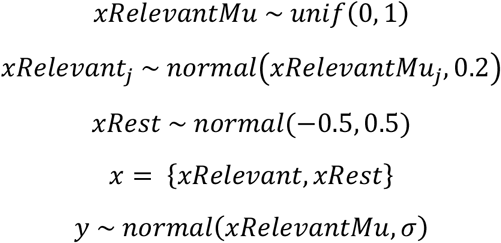

Where ‘unif’ stands for uniform distribution and ‘normal’ stands for normal distribution, j is a site-specific index, and σ is the standard deviation depending on the strength of the climatic effect. We run the model 100 times for each combination of the strength of the climatic effect and the number of sites. We assessed the proportion of models which identified the effect of climate with 95% credible interval of the slope parameter not overlapping zero (for priors and sampling parameters see *Supporting information*).

For a strong climatic effect, 30 sites were enough to identify it in all models. For the medium climatic effect, 30 sites had power greater than 0.95. For the weak climatic effect, 90 sites were necessary to reliably identify the climatic effect (Fig. 3).

**Figure 3:**
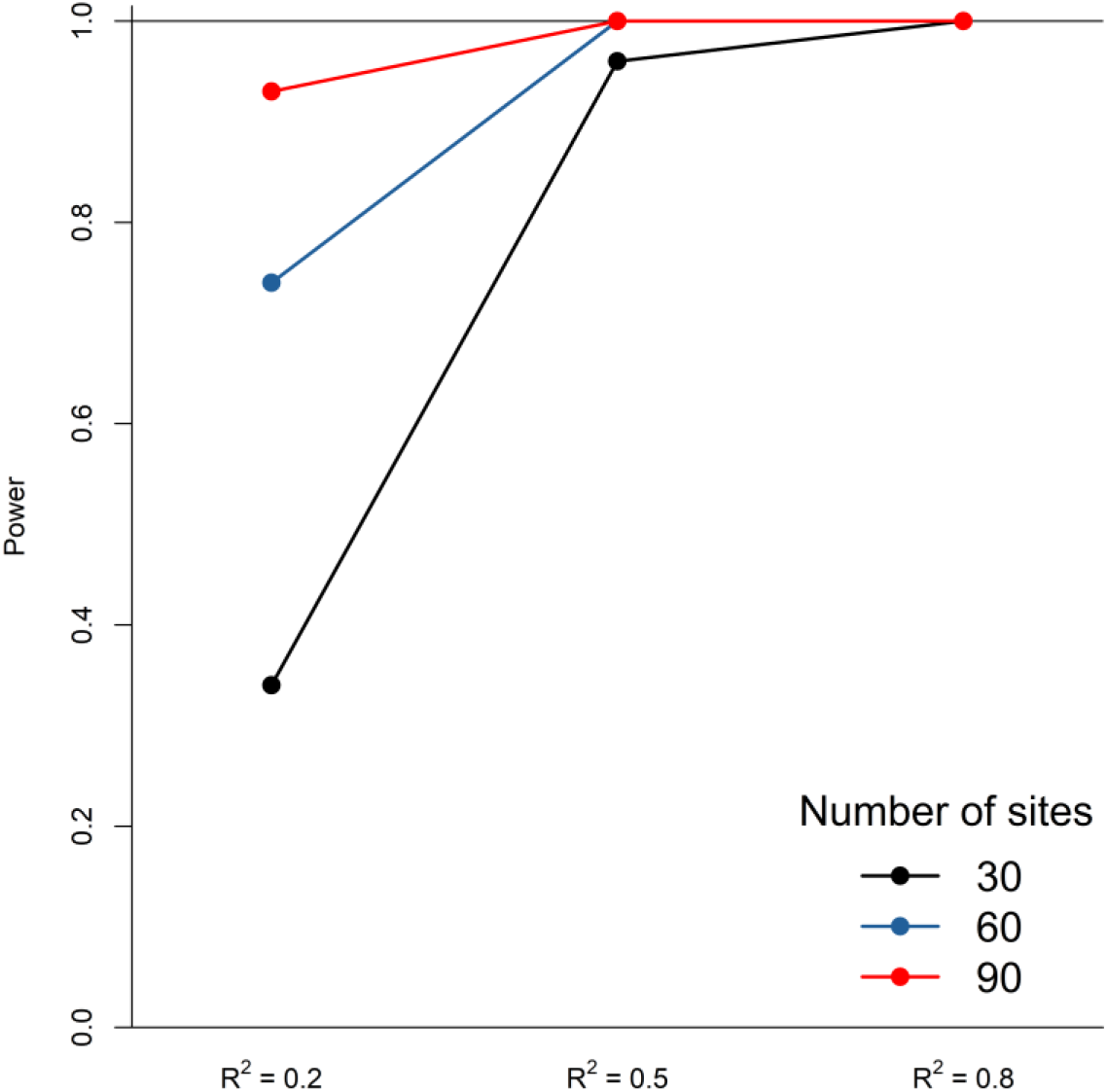
StatClim statistical power based on the number of sites and the strength of the climatic effect expressed as R^2^.

For assessment of false positive rate (type I error), we simulated the response variable as independent of climate. We ran 100 models and the results of all of them were consistent with no effect of climate (95% credible interval of the slope parameter overlapped zero), resulting in a false positive rate < 0.01.

## 3 Case study

To illustrate the potential of StatClim on real data, we analyzed mast seeding of *Fagus sylvatica* and its response to temperature using the dataset previously used to compare other methods for climatic window identification (Cornes et al. 2018; Hacket-Pain et al. 2022; Journé et al. 2025b). Previous studies found that *Fagus sylvatica* L. (European beech) synchronizes its reproduction, and the reproduction is positively affected by summer temperatures of the previous year and negatively by temperature two years before (Nussbaumer et al. 2018; Bogdziewicz et al. 2023).

To account for the specifics of the analyzed dataset, we expanded StatClim to estimate two temperature summaries, included a random effect of population to account for population identity, and autoregressive terms of second order (AR(2)) to account for the periodicity of reproduction of *Fagus sylvatica* (Fig. S2). These modifications also illustrate how StatClim can be extended to account for the complex data (see *Supporting information*).

Consistent with the literature, StatClim identified a negative effect of temperatures two years before the reproduction and a positive effect of temperatures one year before the reproduction (Fig. 4). In addition to what was already known, StatClim results revealed that not average temperatures in identified windows but rather high temperatures play a key role in Fagus reproduction. We also found a negative effect of reproduction in the previous year (the autoregressive term; not two years prior).

**Figure 4:**
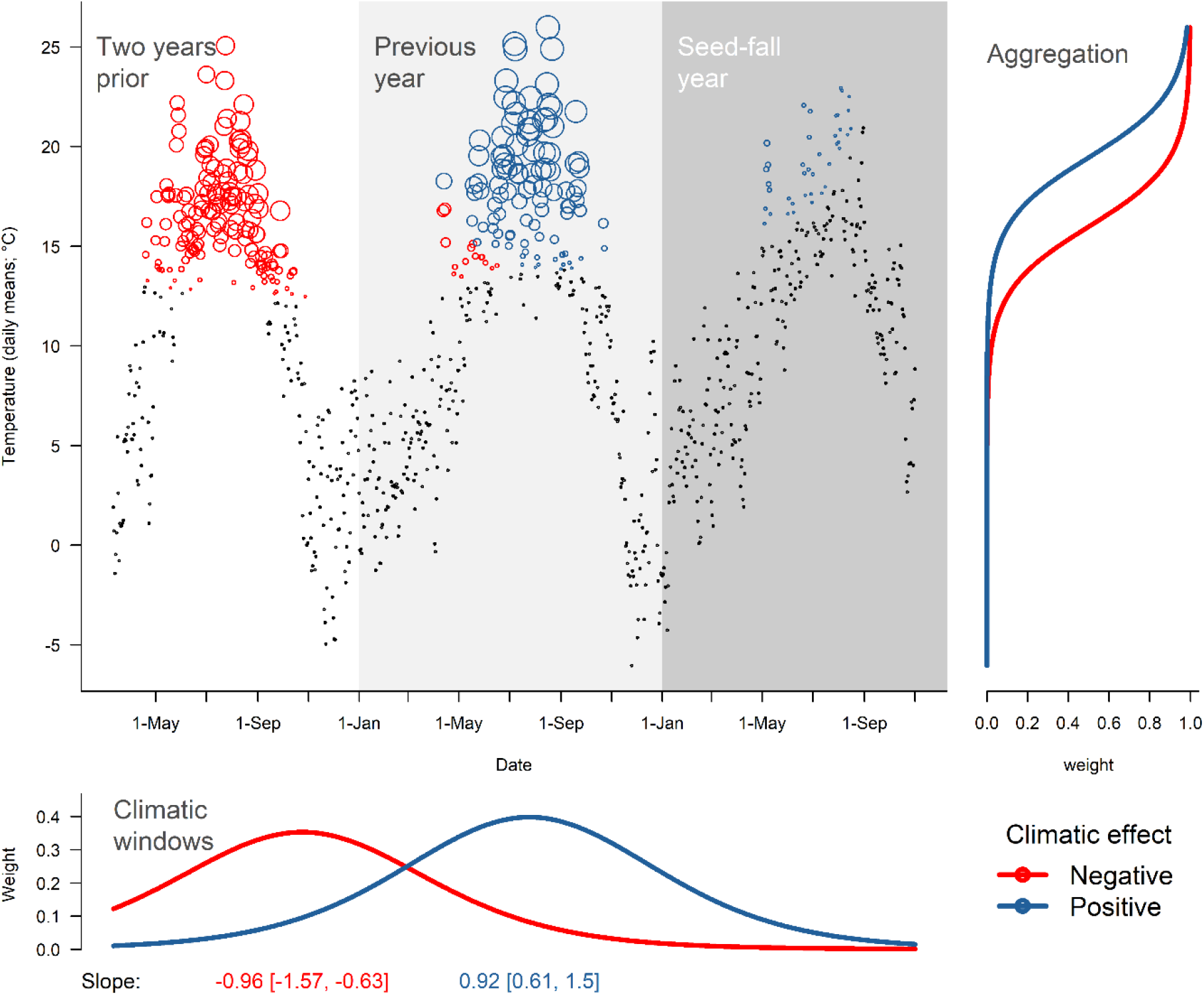
Climatic effects on the reproduction of Fagus sylvatica identified with the StatClim model. The upper left part shows temperature records related to one reproduction event in one population (mast year 1990, population: lon.: 15.11° lat.: 53.15°) with points size corresponding to the maximum assigned weight from the two climatic windows (posterior mean). Temperature records in red contribute primarily (higher weights) to the first climatic window; records in blue contribute to the second climatic window. The bottom part shows the timing of both climatic windows with slope parameters (posterior mean and 95% credible intervals). The right part shows the corresponding aggregations (their logistic windows).

## 4 Discussion

StatClim enables a data-driven summary of climatic information by estimating the climatic window and aggregation, which predicts the response of interest. We presented the model primarily using a space-for-time design but StatClim is also suitable for other designs. Observations of a biological phenomenon replicated in time, such as tree growth increments or plant phenology events (Simmonds et al. 2019), can often be thought of as independent, and thus can be the object of the same type of analyses as space-for-time studies. This is also true for studies replicated in both space and time (e.g., Jarecke et al. 2024; Journé et al. 2025a), together indicating a broad StatClim application potential.

In these fields, StatClim promises an advance in theoretical and applied research, having several advantages compared to its alternatives. Firstly, it identifies not only the relevant climatic window but also the type of aggregation. This differs from previous methods, which are limited to the temporal dimension of the (climatic) data (P-spline regression: Roberts 2008; Peak signal detection: Brakel 2014; Climate sensitivity profile: Thackeray et al. 2016; model selection: van de Pol et al. 2016). This allows StatClim to detect the effects of extreme climatic conditions, which could be overlooked or underestimated in the analysis considering only climatic windows (Körner and Hiltbrunner 2018).

Secondly, StatClim identifies the climatic summary within one model, avoiding an inflated rate of false positives common for the model selection approach (Macphie et al. 2025). Although the false positive rate can be controlled in model selection via randomization (see randwin function in climwin), it is computationally expensive. Furthermore, model selection is often used repeatedly (e.g. for each site separately) leading to an unclear false positive rate (Evers et al. 2021; Anderson-Teixeira et al. 2022). In contrast, StatClim has a very low false positive rate, because it uses zero centered priors (Gelman et al. 2012). Thirdly, the implementation of StatClim within the Bayesian framework provides great flexibility and potential to expand the model (for examples see the *Case study*). This allows users to account for complex data structures often present in ecological datasets while keeping the analysis integrated, which ensures a low false positive rate and propagation of uncertainty. Fourthly, this propagation of uncertainty is important because the climatic summary is estimated with uncertainty, which should be propagated into the uncertainty of slope estimation. Without that, the slope uncertainty can be underestimated, leading again to false positives.

StatClim has its limitations too. First, it requires some knowledge of Bayesian statistics to be adjusted to a specific dataset and its structure, to set appropriate priors, and to fit the model. Fitting complex Bayesian models can be challenging, since the full posterior distribution may have complex geometry (e.g. weakly identified regions or funnels; Neal 2003). This challenge results from the fact that Bayesian inference aims to characterize the full posterior distribution rather than a single optimum (maximum likelihood), which exposes identifiability issues less visible in other approaches. Second, the estimation of complex models can be computationally demanding. While the core StatClim model is relatively quick to estimate, when expanded to include additional features, it can require considerable computational capacities (e.g., hours for the presented *Case study*). Third, the model can be difficult to fit under the presence of multimodality. Multimodality in climate window estimation is very common, since climatic variables are often periodic (e.g., with a period of one day or year). This can lead to multiple climatic windows that predict a response variable similarly well and, consequently, to a problematic estimation of window position and size. Here, data exploration is important, and for that, the model selection approach using the *climwin* package can be very useful. Model selection fits all the possible models and thus explores any multimodality which can then be accounted for by expanding the core StatClim model, for example, by including an extra climatic window.

For StatClim applications, we provide the *Stan* code of several model variants (https://github.com/adamklimes/StatClim) as a starting point:

1. core StatClim model, which estimates one climatic window and corresponding aggregation.
2. expanded StatClim model which estimates two climatic windows and their aggregation, one random effect (used in the case study to account for population identity), and autoregressive terms of second order (AR(2)); used in the presented case study.
3. separated StatClim submodels estimating either only the climatic window (and using mean value within this window) or only the most informative aggregation across fixed time window. These submodels can be useful for model development, analysis workflow (Gelman et al. 2020), or hypotheses which concern only the climatic window or aggregation.
4. expanded StatClim with multiple (micro)climatic variables (with the climatic window and aggregation estimated for each of them). In this model, the prior for slope parameters can be defined as zero centered with a normal or Laplace distribution to implement the Bayesian version of the ridge or LASSO regression respectively (Park and Casella 2008; with ridge or LASSO penalty either fixed or estimated; Liao 2023). Ridge or LASSO regression can be especially useful when the goal of the analysis is to identify the subset of (micro)climatic variables with an effect on the variable of interest, because they effectively provide variable selection and regularization improving prediction power and interpretability (Tibshirani 1996).
5. StatClim using Weibull function instead of a logistic window for climate window estimation (aggregation is still done by logistic window). The Weibull function can be used to assign different weights to climatic records, for example, modelling a diminishing effect of a (micro)climatic variable with increasing time difference from an event of interest, similarly to the climwin function weightwin (Bailey and van de Pol 2016).

These different models illustrate that the framework implemented in StatClim is general and the core StatClim model is a building block which can be expanded or incorporated into other models as needed for a particular analysis. StatClim can be also accompanied by other climate window estimation methods, which can provide efficient data exploration or robustness check of results. In summary, StatClim substantially broadens options for (micro)climate time series summary estimation and thus contributes to the data-driven ecological research.

## Authors Contributions

Adam Klimeš: conceptualization, methodology, software, writing – original draft, writing – review & editing. Martin Macek: writing – review & editing. Jan Wild: writing – review & editing. Josef Brůna: project administration, writing – review & editing. Martin Kopecký: supervision, writing – review & editing.

## Acknowledgements

The authors are grateful to members of the Geoecology Department of the Institute of Botany for their comments on early versions of the manuscript.

## Data availability statement

All data used are publicly available. The code for all analyzes is available here: https://github.com/adamklimes/StatClim.

## References

Akaike H (1974) A new look at the statistical model identification. IEEE Trans Automat Contr 19:716–723. 10.1109/TAC.1974.1100705

Anderson-Teixeira KJ, Herrmann V, Rollinson CR, et al (2022) Joint effects of climate, tree size, and year on annual tree growth derived from tree-ring records of ten globally distributed forests. Glob Chang Biol 28:245–266. 10.1111/gcb.15934

Bailey LD, van de Pol M (2016) climwin: An R Toolbox for Climate Window Analysis. PLoS One e0167980:. 10.1371/journal.pone.0167980

Bogdziewicz M, Journé V, Hacket-Pain A, Szymkowiak J (2023) Mechanisms driving interspecific variation in regional synchrony of trees reproduction. Ecol Lett 26:754–764. 10.1111/ele.14187

Brakel JPG van (2014) Robust peak detection algorithm using z-scores. In: Stack Overflow. https://stackoverflow.com/questions/22583391/peak-signal-detection-in-realtime-timeseries-data/22640362#22640362

Carpenter B, Gelman A, Hoffman MD, et al (2017) Stan : A probabilistic programming language. J Stat Softw 76:1–32. 10.18637/jss.v076.i01

Cornes RC, van der Schrier G, van den Besselaar EJM, Jones PD (2018) An Ensemble Version of the E-OBS Temperature and Precipitation Data Sets. Journal of Geophysical Research: Atmospheres 123:9391–9409. 10.1029/2017JD028200

Currier CM, Sala OE (2022) Precipitation versus temperature as phenology controls in drylands. Ecology 103:. 10.1002/ecy.3793

Evers SM, Knight TM, Inouye DW, et al (2021) Lagged and dormant season climate better predict plant vital rates than climate during the growing season. Glob Chang Biol 27:1927–1941. 10.1111/gcb.15519

Gelman A, Hill J, Yajima M (2012) Why We (Usually) Don’t Have to Worry About Multiple Comparisons. J Res Educ Eff 5:189–211. 10.1080/19345747.2011.618213

Gelman A, Vehtari A, Simpson D, et al (2020) Bayesian Workflow. arXiv preprint arXiv:2011:

Glądalski M, Bańbura M, Kaliński A, et al (2018) Hatching delays in great tits and blue tits in response to an extreme cold spell: a long-term study. Int J Biometeorol 62:1437–1445. 10.1007/s00484-018-1541-3

Hacket-Pain A, Foest JJ, Pearse IS, et al (2022) MASTREE+: Time-series of plant reproductive effort from six continents. Glob Chang Biol 28:3066–3082. 10.1111/gcb.16130

Jarecke KM, Bladon KD, Meinzer FC, Wondzell SM (2024) Impact of rainfall and vapor pressure deficit on latewood growth and water stress in Douglas-fir in a Mediterranean climate. For Ecol Manage 551:. 10.1016/j.foreco.2023.121529

Journé V, Kelly D, Hacket-Pain A, et al (2025a) Weather drivers of reproductive variability in perennial plants and their implications for climate change risks. Nature Communications 16:. 10.1038/s41467-025-64300-6

Journé V, Simmonds EG, Barczyk MK, Bogdziewicz M (2025b) Comparing statistical methods for detecting weather cues of mast seeding in European beech (Fagus sylvatica) across Europe. Agric For Meteorol 375:. 10.1016/j.agrformet.2025.110857

Kharouba HM, Williams JL (2024) Forecasting species’ responses to climate change using space-for-time substitution. Trends Ecol Evol 39:716–725. 10.1016/j.tree.2024.03.009

Klinges DH, Baecher JA, Lembrechts JJ, et al (2024) Proximal microclimate: Moving beyond spatiotemporal resolution improves ecological predictions. Global Ecology and Biogeography 33:. 10.1111/geb.13884

Körner C, Hiltbrunner E (2018) The 90 ways to describe plant temperature. Perspect Plant Ecol Evol Syst 30:16–21. 10.1016/j.ppees.2017.04.004

Lenarz MS, Nelson ME, Schrage MW, Edwards AJ (2009) Temperature Mediated Moose Survival in Northeastern Minnesota. J Wildl Manage 73:503–510. 10.2193/2008-265

Liao J (2023) A new Bayesian lasso and ridge regression with a practically meaningful parameterization and a simple weakly informative prior. Stat 12:1–9. 10.1002/sta4.540

Lovell RSL, Collins S, Martin SH, et al (2023) Space-for-time substitutions in climate change ecology and evolution. Biological Reviews 98:2243–2270. 10.1111/brv.13004

Macek M, Kopecký M, Wild J (2019) Maximum air temperature controlled by landscape topography affects plant species composition in temperate forests. Landsc Ecol 34:2541–2556. 10.1007/s10980-019-00903-x

Macphie KH, Morris S, Morris A, et al (2025) Addressing biases in sliding window analysis gives new insight into the response of parturition date to weather in a wild mammal. Oikos e11372:. 10.1002/oik.11372

McCowan LSC, Griffith SC (2021) Baked eggs: catastrophic heatwave-induced reproductive failure in the desert-adapted Zebra Finch (Taeniopygia guttata). Ibis 163:1207–1216. 10.1111/ibi.12958

Moreno-Rueda G, Pizarro M (2008) Temperature differentially mediates species richness of birds of different biogeographic types. Ardea 96:115–120. 10.5253/078.096.0113

Neal RM (2003) Slice sampling. Ann Stat 31:743–748. 10.2307/3448415

Nussbaumer A, Waldner P, Apuhtin V, et al (2018) Impact of weather cues and resource dynamics on mast occurrence in the main forest tree species in Europe. For Ecol Manage 429:336–350. 10.1016/j.foreco.2018.07.011

Park T, Casella G (2008) The Bayesian Lasso. J Am Stat Assoc 103:681–686. 10.1198/016214508000000337

Roberts AMI (2008) Exploring relationships between phenological and weather data using smoothing. Int J Biometeorol 52:463–470. 10.1007/s00484-007-0141-4

Rubio-Cuadrado Á, Camarero JJ, Bosela M (2022) Applying climwin to dendrochronology: A breakthrough in the analyses of tree responses to environmental variability. Dendrochronologia (Verona) 71:. 10.1016/j.dendro.2021.125916

Simmonds EG, Cole EF, Sheldon BC (2019) Cue identification in phenology: A case study of the predictive performance of current statistical tools. Journal of Animal Ecology 88:1428–1440. 10.1111/1365-2656.13038

Sollmann R (2024) Estimating the temporal scale of lagged responses in species abundance and occurrence. Ecosphere 15:. 10.1002/ecs2.4704

Svenning JC, Sandel B (2013) Disequilibrium vegetation dynamics under future climate change. Am J Bot 100:1266–1286. 10.3732/ajb.1200469

Thackeray SJ, Henrys PA, Hemming D, et al (2016) Phenological sensitivity to climate across taxa and trophic levels. Nature 535:241–245. 10.1038/nature18608

Tibshirani R (1996) Regression Shrinkage and Selection via the Lasso. Journal of the Royal Statistical Society: Series B (Methodological) 58:267–288. 10.1111/j.2517-6161.1996.tb02080.x

van de Pol M, Bailey LD, McLean N, et al (2016) Identifying the best climatic predictors in ecology and evolution. Methods Ecol Evol 7:1246–1257. 10.1111/2041-210X.12590

van de Pol M, Cockburn A (2011) Identifying the critical climatic time window that affects trait expression. American Naturalist 177:698–707. 10.1086/659101

Van Kampen R, Fisichelli N, Zhang YJ, Wason J (2022) Drought timing and species growth phenology determine intra-annual recovery of tree height and diameter growth. AoB Plants 14:. 10.1093/aobpla/plac012

Wyver C, Potts SG, Edwards M, et al (2023) Spatio-temporal shifts in British wild bees in response to changing climate. Ecol Evol 13:. 10.1002/ece3.10705

