## Supporting information for "Data-driven identification of the ecologically relevant climatic windows and climate summaries with StatClim model"

Content:

- Sampling and prior specification
- Case study details
- Figure S1
- Figure S2
- References

**Sampling and prior specification**

We used in the power analysis and the case study (and StatClim outputs section) zero centered normally distributed priors with standard deviation 2 for all parameters (half-normally distribution for variance parameters) apart from start and width of climatic window where we used Dirichlet distribution with parameter values 1.5. We fitted every model using 4 chains with 2000 iterations each, half of them as a warm-up phase. Shape parameter was 0.1 for sub-model 1 and 0.05 for sub-model 2. For code see link in the Data Availability Statement.

**Case study details**

In the Case study, we reanalyzed a subset of a dataset from Journe et al. (2025) coming originally from MASTREE+ database (Hacket-Pain et al. 2022) and the E-OBS dataset (Cornes et al. 2018) accessed at https://github.com/ValentinJourne/weatheRcues/tree/main/Application_MASTREE.

We selected 11 Fagus sylvatica populations in Poland with continuous records of the percentage of reproducing individuals since 1990 (up until 2019; Fig. S2); they represent a homogeneous set in terms of data collection methods and units used. Following Journé et al. (2025), the response variable was the percentage of reproducing individuals + 1 transformed using natural logarithm and temperature measurements were daily mean temperatures for 966 days ending on the first of November of the year when the reproduction was recorded (here we added one more year to be able to identify also the effect of temperature two years before the reproduction). To account for population identity, we added a random effect of population and to account for the periodicity of reproduction of Fagus sylvatica, which is 2 or 3 years (Fig. S2), we added autoregressive terms of second order (AR(2); i.e., reproduction one and two years before).

Since previous results suggested that temperatures both one year and two years before the reproduction have an effect and these effects are different (Nussbaumer et al. 2018; positive and negative, respectively; Bogdziewicz et al. 2023), we estimated not one but two temperature summaries. For each summary, we estimated the beginning of the time window and its width, aggregation type, and the effect of the summary on reproduction (slope).

We used the same specification of priors and sampling parameters as for the power analysis.

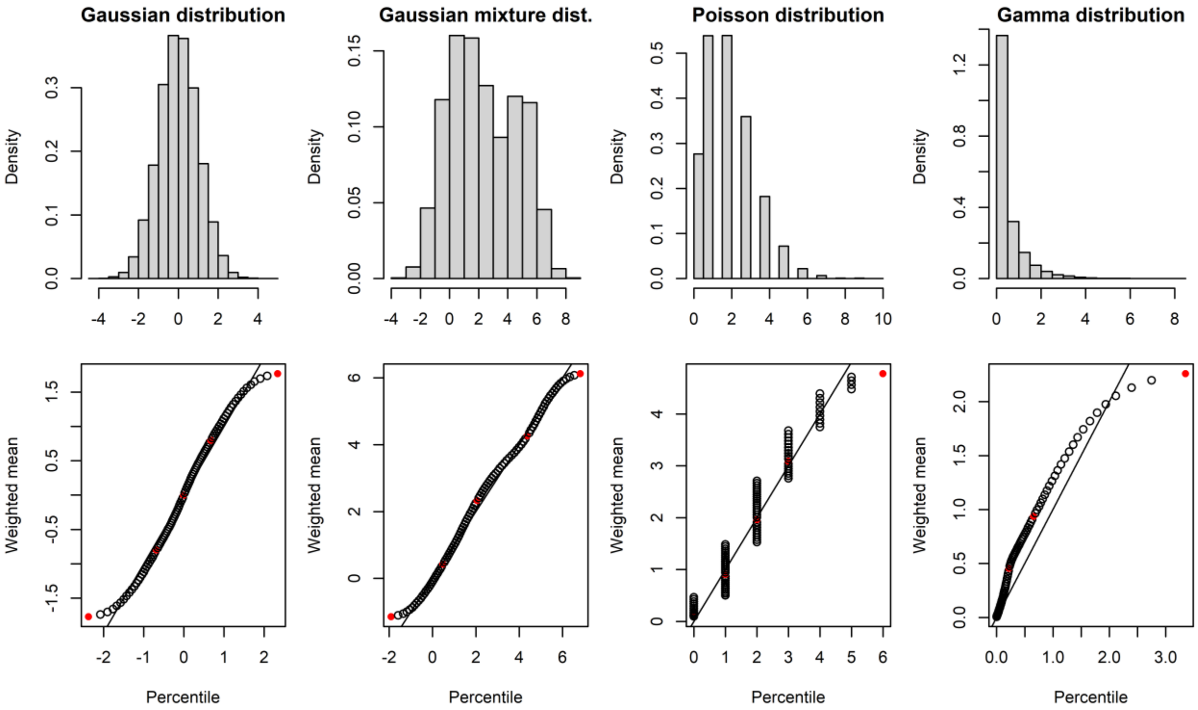


***Figure S1****: Comparison of percentile and weighted mean with weights based on a logistic window for four different distributions. Red points highlight 1st, 25th, 50th, 75th, and 99th percentile; shape parameter (see the main text) was set to 0.05. Values of weighted mean are very similar to percentile for all four distributions thus summarization based on weighted mean can be interpreted similarly to summarization based on percentile.*

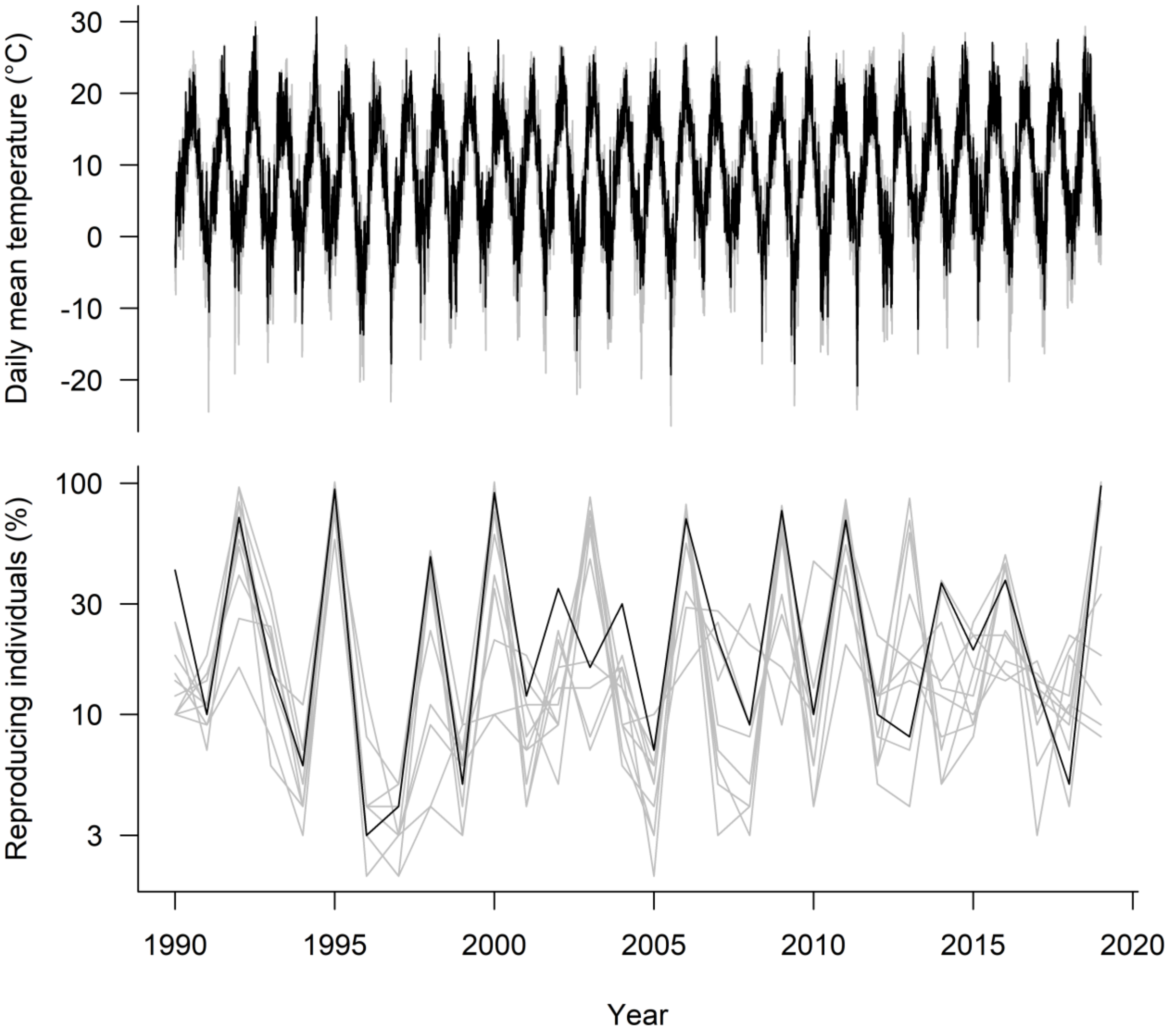


***Figure S2****: Temperature and reproduction (+1 and log-scale) of 11 populations of Fagus sylvatica. One population is highlighted in black (population: lon.: 15.11° lat.: 53.15°). Data from Journé et al. (2025).*

**References**


Bogdziewicz M, Journé V, Hacket-Pain A, Szymkowiak J (2023) Mechanisms driving interspecific variation in regional synchrony of trees reproduction. Ecol Lett 26:754–764. https://doi.org/10.1111/ele.14187

Cornes RC, van der Schrier G, van den Besselaar EJM, Jones PD (2018) An Ensemble Version of the E-OBS Temperature and Precipitation Data Sets. Journal of Geophysical Research: Atmospheres 123:9391–9409. https://doi.org/10.1029/2017JD028200

Hacket-Pain A, Foest JJ, Pearse IS, et al (2022) MASTREE+: Time-series of plant reproductive effort from six continents. Glob Chang Biol 28:3066–3082. https://doi.org/10.1111/gcb.16130

Journé V, Simmonds EG, Barczyk MK, Bogdziewicz M (2025) Comparing statistical methods for detecting weather cues of mast seeding in European beech (Fagus sylvatica) across Europe. Agric For Meteorol 375:. https://doi.org/10.1016/j.agrformet.2025.110857

Nussbaumer A, Waldner P, Apuhtin V, et al (2018) Impact of weather cues and resource dynamics on mast occurrence in the main forest tree species in Europe. For Ecol Manage 429:336–350. https://doi.org/10.1016/j.foreco.2018.07.011
